# Genome-wide maps of transcription factor footprints identify noncoding variants rewiring gene regulatory networks

**DOI:** 10.64898/2026.03.03.708783

**Authors:** Jiecong Lin, Wenyang Dong, Jiaxiang Zhang, Chen Xie, Xiaoxi Jing, Junpeng Zhao, Kaiyan Ma, Hongen Kang, Yilin Jiang, Xiaoliang Sunney Xie, Yajie Zhao

## Abstract

Genome-wide association studies have identified millions of noncoding loci linked to human traits, yet how these variants alter gene regulation remains a major challenge, particularly for rare variants where whole-genome sequencing cohorts and high-resolution functional annotations remain limited. Here we show that single-molecule deaminase footprinting (FOODIE) in K562 cells captures up to 103-fold heritability enrichment for erythroid traits despite covering 0.12% of the genome. We introduce varTFBridge, integrating FOODIE footprinting with AlphaGenome variant effect prediction to identify causal noncoding variants altering transcription factor (TF)-mediated regulation. Applied to 490,640 UK Biobank genomes across 13 erythrocyte traits, varTFBridge prioritises 113 high-confidence regulatory variants (104 common, 9 rare), encompassing 2,173 linkages along the variant–TF binding– gene–trait cascade across 64 TFs and 108 genes. varTFBridge recapitulates rs112233623 and resolves its mechanism: GATA1/TAL1 co-binding disruption at a *CCND3* enhancer altering red blood cell count and volume.

## Introduction

Genome-wide association studies (GWAS) have identified millions of trait-associated variants, yet most reside in noncoding regions and remain functionally uncharacterized^1–4^. Fine-mapping has improved the localization of causal variants, but the underlying regulatory mechanisms are often unresolved^5–9^. Moreover, rare variants, which can pinpoint causal genes with large effect sizes^10^, have been studied primarily in coding regions owing to the difficulty of interpreting noncoding effects^11^. Characterizing noncoding variants and establishing functional links between these loci, their target genes, and the regulatory landscape thus remains a central challenge.

Transcription factors (TFs) bind specific noncoding genomic regions to regulate gene expression^12,13^, and variants within TF binding motifs can disrupt or create binding sites, thereby altering target gene expression and contributing to disease^14–16^. Understanding these mechanisms has direct therapeutic relevance: the gene-editing therapy Casgevy^17^ targets a BCL11A enhancer identified through GWAS to reactivate fetal hemoglobin for the treatment of β-thalassemia^18–20^. The release of whole-genome sequencing (WGS) data from nearly 500,000 UK Biobank participants^21^, encompassing approximately 1.5 billion variants, now provides an opportunity to systematically investigate how noncoding regulatory variants modulate disease risk.

Recently, single-molecule deaminase footprinting (FOODIE) has enabled genome-wide measurement of TF binding at near-single-base resolution^22^ providing direct, cell-type-specific maps of TF occupancy that can pinpoint the exact binding sites affected by noncoding variants^23^. Concurrently, AlphaGenome^24,25^, a deep-learning model trained on thousands of epigenomic profiles, has demonstrated strong performance in predicting how single-nucleotide variants alter chromatin accessibility, histone modifications, and TF binding across cell types^26^. Together, FOODIE’s ability to resolve TF binding at base-pair resolution and AlphaGenome’s capacity to predict variant effects across multiple epigenomic layers offer complementary strengths for decoding the regulatory mechanisms of noncoding variants. However, no integrative framework currently combines these approaches.

Here we present varTFBridge, a framework that integrates high-resolution FOODIE footprinting data with AlphaGenome variant effect prediction to prioritize causal noncoding variants across the allele frequency spectrum. The framework identifies trait-associated variants within TF footprints through GWAS fine-mapping for common variants and, for the first time, footprint-wide burden testing for rare variants, then maps their functional consequences using motif-based binding prediction, TF footprint–gene linking, and Alphagenome variant effect scoring. We first show that FOODIE footprints outperform DNase-seq and ATAC-seq in TF binding measurement, enhancer–gene linkage prediction, and erythrocyte trait heritability enrichment in K562 cells. FOODIE libraries were generated using an updated protocol (Vazyme Hyperactive FOODIE Library Prep Kit) with improved efficiency. We then apply varTFBridge to 490,640 UK Biobank genomes across 13 erythrocyte traits, mapping 209 common (MAF ≥ 0.1%) and 18 rare (MAF < 0.1%) variants to TF binding sites and target genes. Convergent-evidence filtering prioritises 113 high-confidence variants, encompassing 2,173 unique variant–TF binding–gene–trait linkages. The framework recapitulates the causal variant rs112233623 and resolves its molecular mechanism: disruption of a GATA1/TAL1 co-binding motif at a CCND3 enhancer, resulting in higher mean corpuscular volume and lower red blood cell count.

## Results

### The varTFBridge framework

We developed varTFBridge, an integrated workflow that uses whole-genome sequencing data from 490,640 UK Biobank individuals^21^ to identify TF-mediated noncoding regulatory variants for blood traits in a cell-type-specific context. The workflow proceeds in two stages: (1) identification of trait-associated common and rare variants, and (2) functional dissection of putative causal variants at the level of TF binding sites and target genes.

In the variant association stage (Fig. 1c–d), we performed GWAS followed by genome-wide fine-mapping (GWFM) with SBayesRC^27^ for common variants (MAF ≥ 0.1%) to pinpoint putative causal variants within cell-type-specific TF footprints (Methods). For rare variants (MAF < 0.1%), we conducted footprint-based burden tests by collapsing variants within each footprint, followed by leave-one-variant-out analysis^28^ to isolate the specific driver variant within each significant footprint (Methods).

**Fig. 1.**
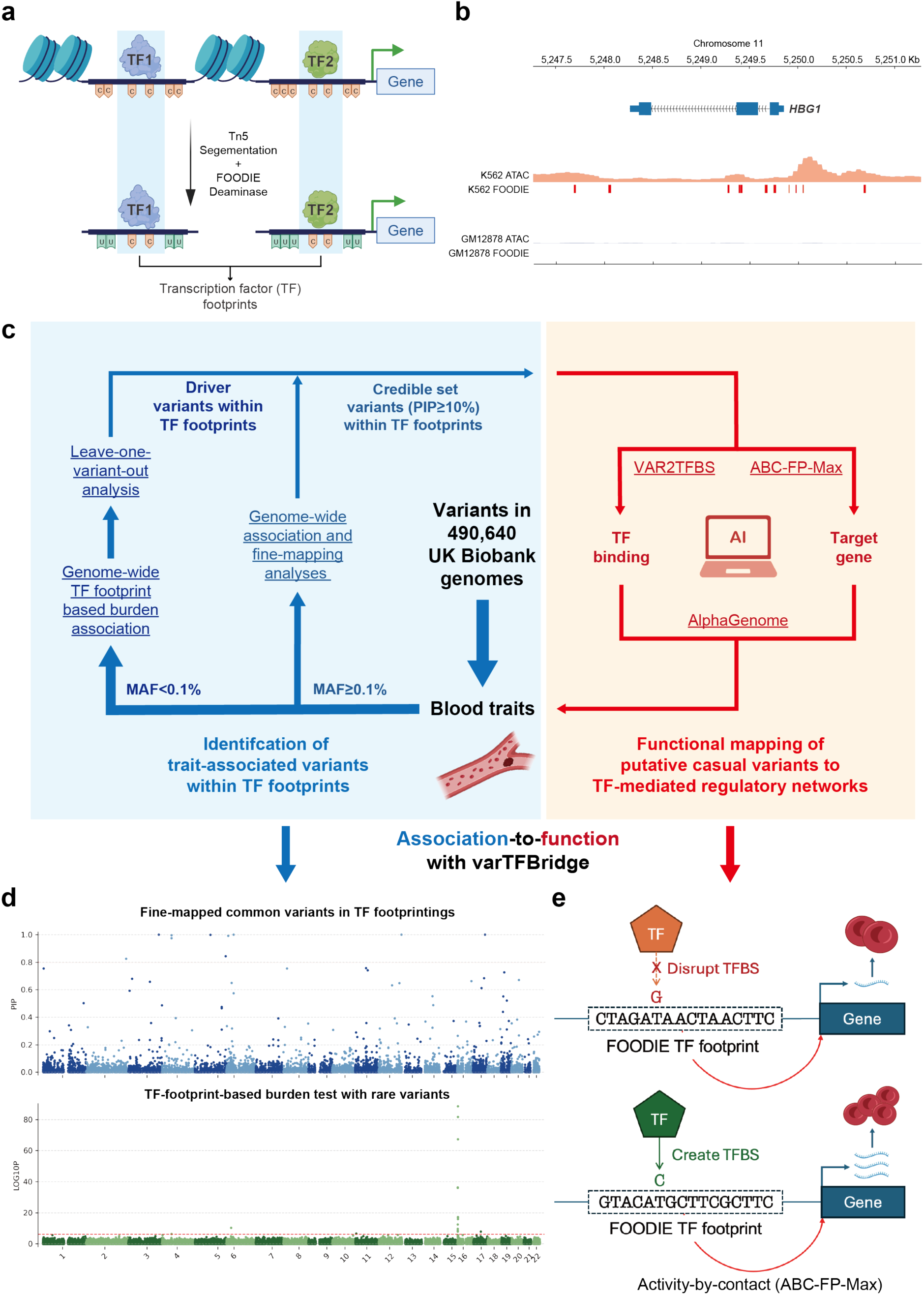
Overview of varTFBridge and FOODIE footprinting. **a,** Schematic of FOODIE, in which nuclei are treated with a cytosine deaminase following Tn5 transposition, converting cytosines to uracils in proportion to chromatin accessibility. **b**, Genome browser view of the *HBG1* locus encoding the γ-globin chain and highly expressed in K562. ATAC-seq tracks for K562 and GM12878 cells (rows 1 and 3) are shown alongside corresponding FOODIE footprint tracks (rows 2 and 4). **c**, The varTFBridge workflow. Procedures highlighted in blue identify blood-trait–associated credible variants from 490,640 UK Biobank participants, whereas procedures in red functionally map these variants to transcription factor–mediated gene regulatory networks. Underlined text denotes methods implemented within the varTFBridge framework. Arrows denote the progression from genetic association to functional interpretation. **d**, Manhattan plots for fine-mapped common variants overlapping FOODIE footprints (top) and footprint-based rare-variant burden tests (bottom) associated with blood cell traits. Posterior inclusion probabilities (PIPs) for common variants were estimated using genome-wide fine-mapping with SBayesRC. Each point represents a variant (top) or a FOODIE footprint (bottom) plotted by genomic position. **e**, Conceptual illustration of how regulatory variants that alter transcription factor binding, either disrupting or creating binding sites, to modulate gene expression and drive red blood cell phenotypic variation. TFBS: transcription factor binding site.

In the functional dissection stage, we used FIMO^29^ to scan JASPAR^30^ TF binding motifs within footprints and assessed how putative causal variants affect predicted binding affinity (termed VAR2TFBS; Methods). We adapted the ABC-Max model^8,31^ by integrating TF footprints (termed ABC-FP-Max; Methods) to link variants to target genes (Fig. 1e). Finally, we employed AlphaGenome to predict variant effects across multiple epigenomic layers, including histone modifications, TF binding, and chromatin accessibility (Methods). We then integrated these predictions through convergent-evidence filtering, requiring concordant support from motif disruption, TF expression, and variant-to-gene linkage, to produce a high-confidence variant–TF binding–gene–trait regulatory resource (Methods). We applied this framework to 13 erythroid traits in K562 cells.

### FOODIE footprints improve TF binding measurement and enhancer–gene prediction

We performed FOODIE single-molecule footprinting in K562 and GM12878 cells using an updated kit (Vazyme TD721), generating genome-wide TF footprint profiles at near-single-base resolution (Fig. 1a–b). To evaluate FOODIE’s capacity to capture TF binding alongside existing assays, we applied the footprint-calling protocol to K562 FOODIE data (Methods), yielding 188,484 footprints with a median length of 19 bp. Comparison with DNase and ATAC-seq data in K562 from ENCODE showed that 91.8% of footprints overlap at least one chromatin accessibility peak and 61% are supported by both, consistent with TF footprints predominantly residing in open chromatin (Fig. 2a).

**Fig. 2.**
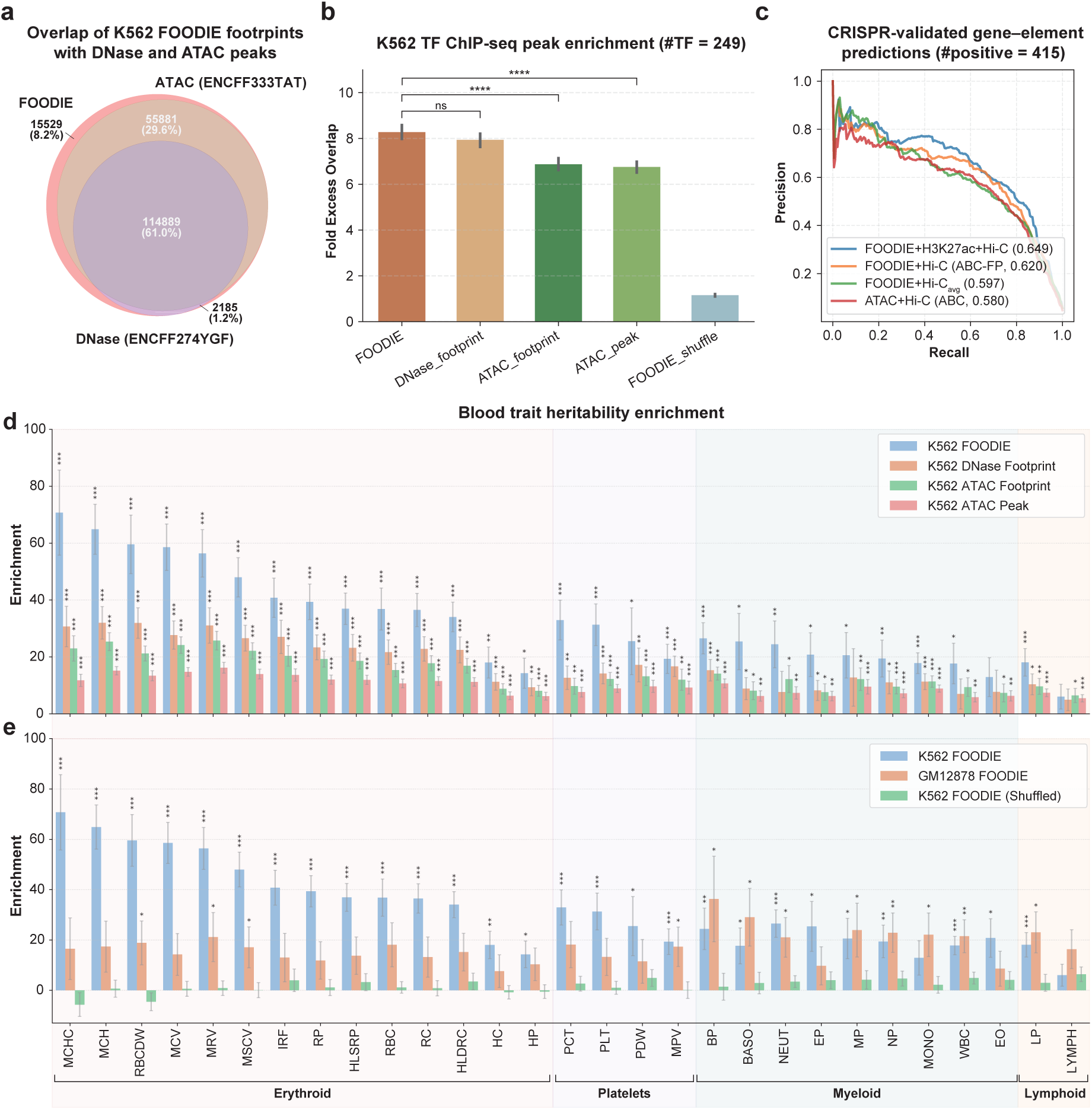
Genome-wide characterization of FOODIE transcription factor footprints. **a.** Venn diagram showing overlap between FOODIE footprints and DNase-seq and ATAC-seq peaks in K562 cells. ENCODE sample identifiers are provided in parentheses. **b.** Comparison of transcription factor (TF) footprints and open chromatin profiles based on fold excess overlap with TF ChIP–seq peaks. Fold excess overlap with TF ChIP–seq peaks in K562 cells is shown for FOODIE, ATAC-seq and DNase-seq footprints, alongside ATAC-seq peaks and shuffled FOODIE footprints (317 ChIP–seq experiments covering 249 unique TFs). Error bars denote one standard error. **** indicates *P* < 1 × 10^−4^ by two-sided *t*-test; ns denotes non-significant differences (*P* > 0.05). **c.** Precision–recall curves for classifiers predicting enhancer–gene (EG) pairs. Positive EG pairs are defined as those in which CRISPR perturbation of a distal element significantly decreases expression of the target gene. In total, 415 positive EG pairs among 7,238 tested EGs were used for benchmarking. Curves show performance across different combinations of epigenomic profiles within the Activity-by-Contact (ABC) framework. ABC-FP denotes the footprint–gene linkage score proposed in this study, whereas ABC denotes the original ATAC–based ABC score. **d.** Bar plot showing heritability enrichment (proportion of heritability divided by the proportion of SNPs) for four footprint annotations and a published ENCODE ATAC-seq peak annotation in K562 cells across 29 blood cell–related traits. *** indicates *P* < 1 × 10^−3^by one-sided *t*-test; ** indicates *P* < 1 × 10^−2^by one-sided *t*-test; * indicates *P* < 5 × 10^−2^by one-sided *t*-test. **e.** Bar plot of heritability enrichment for FOODIE footprints in K562 and GM12878 cells across 29 blood cell–related traits. Shuffled K562 FOODIE footprints serve as a baseline. Traits are color-coded by the respective lineage. Blood cell traits indicated in the figure are as follow: RC: Reticulocyte count; IRF: Immature fraction of reticulocytes; HP: Hematocrit; HC: Hemoglobin concentration; HLDRC: High light scatter reticulocyte count; MCHC: Mean corpuscular hemoglobin concentration; MCH: Mean corpuscular hemoglobin; HLSRP: High light scatter reticulocyte percentage; RBCDW: Red cell distribution width; MCV: Mean corpuscular volume; MRV: Mean reticulocyte volume; MSCV: Mean sphered corpuscular volume; RP: Reticulocyte fraction of red cells; RBC: Red blood cell count; LP: Lymphocyte percentage of white cells; LYMPH: Lymphocyte count; Monocyte: Monocyte count; NP: Neutrophil percentage of white cells; NEUT: Neutrophil count; EP: Eosinophil percentage of white cells; BASO: Basophil count; WBC: White blood cell count; MP: Monocyte percentage of white cells; EO: Eosinophil count; BP: Basophil percentage of white cells. PLT: Platelet count; MPV: Mean platelet volume; PCT: Plateletcrit; PDW: Platelet distribution width. MONO, NP, NEUT, EP, BASO, WBC, MP, EO, and BP are non-erythroid myeloid traits; LP and LYMPH are lymphoid traits; PLT, MPV, PDW, and PCT are megakaryocytic traits; and RC, IRF, HP, HC, HLDRC, MCHC, MCH, HLSRP, RBCDW, MCV, MRV, MSCV, RP, and RBC are erythroid traits.

We compared FOODIE, DNase, and ATAC-derived footprints against K562 ChIP-seq data (249 TFs; 317 ENCODE experiments^32^) using fold excess overlap^14^ (Fig. 2b). DNase and ATAC footprints were called using footprint-tools^12^ and HINT^33^, respectively. FOODIE footprints exhibited the highest excess overlap with ChIP-seq (8.3×, SD 2.2), followed by DNase (7.9×, SD 2.1), ATAC footprints (6.8×, SD 1.7), and ATAC peaks (6.8×, SD 1.7), confirming that FOODIE more accurately captures cell-type-specific TF binding.

To link variants within TF footprints to target genes, we adapted the Activity-by-Contact (ABC) framework^31^ to develop ABC-FP. Because individual FOODIE footprints are narrow (median ∼15 bp), we extended them by 50 bp and merged nearby footprints to form 70,670 regulatory elements (median length 534 bp) suitable for element-level activity quantification. Following the ABC protocol^34^, we calculated element activity (ATAC-seq and H3K27ac ChIP-seq) and Hi-C contact frequency to all genes within 5 Mb, weighting each footprint’s regulatory effect by its chromatin contact frequency with target gene promoters (Methods).

We benchmarked ABC-FP against a K562 CRISPR perturbation dataset^35,36^ containing 7,238 element–gene pairs and 418 validated positives. FOODIE footprints were enriched within CRISPR-validated enhancers (odds ratio 3.51; Supplementary Figure 1). ABC-FP with ATAC, H3K27ac, and K562-specific Hi-C achieved the highest accuracy (AUPRC = 0.649; Fig. 2c). Because cell-type-specific Hi-C data are unavailable for most cell types, we also tested a cell-type-agnostic reference Hi-C map^8^; ABC-FP still achieved AUPRC = 0.597, surpassing the original ATAC-based ABC score (0.58).

These results demonstrate that FOODIE footprints outperform ATAC-seq in capturing TF binding and predicting enhancer–gene interactions. However, ChIP-seq peaks are not a true gold standard for TF binding^37^, as peak regions (200–500 bp) exceed canonical footprints (6–20 bp)^14^, potentially underestimating enrichment. We therefore turned to heritability analyses to assess whether FOODIE footprints are enriched for trait-relevant genetic variation.

### FOODIE footprints are disproportionately enriched in the heritability of erythrocyte traits

Given the erythroid differentiation potential of K562 cells^38^, we examined heritability enrichment of K562 FOODIE footprints across 28 UK Biobank blood traits (13 erythroid, 4 platelet, 9 myeloid, 2 lymphoid) using stratified LD score regression (S-LDSC) with the baseline-LD model v2.0^39^ (Methods). We benchmarked FOODIE against DNase-seq and ATAC-seq footprints, as well as ATAC-seq peaks. Following previous study^14^, we extended TF footprint annotations by 50 bp to ensure sufficient SNP coverage for robust S-LDSC estimation; ATAC-seq peaks were used without extension.

K562 FOODIE footprints yielded the highest heritability enrichment in 28 of 29 blood traits, significantly exceeding ATAC and DNase footprints in erythroid and myeloid traits (Fig. 2d). FOODIE showed median 40-fold and 21-fold enrichment for erythroid and myeloid traits, respectively, compared with 12-fold for lymphoid traits, consistent with the erythroid identity of K562. FOODIE footprint regions comprise 0.52% of the genome yet exhibit 71-fold enrichment for mean corpuscular hemoglobin concentration (MCHC), explaining 40.27% of the trait’s heritability, compared with 31-fold (DNase footprint) and 23-fold (ATAC footprint) (Supplementary Data 6).

Notably, without this extension, raw FOODIE footprints, covering 0.12% of the genome, achieved even higher peak enrichment for most erythroid traits (e.g., ∼103-fold for MCHC compared with ∼71-fold after extension), but enrichment for non-erythroid traits, particularly myeloid (1 of 9 significant) and lymphoid (0 of 2 significant), largely failed to reach significance (Supplementary Fig. 3 and Supplementary Data 6). The 50 bp extension thus captures flanking regulatory context that broadens the signal across blood cell lineages while moderately attenuating peak erythroid enrichment.

To test lineage specificity, we generated GM12878 FOODIE footprints and performed S-LDSC alongside K562 (Fig. 2e). GM12878 footprints showed higher enrichment for lymphoid traits and 6 of 9 myeloid traits, particularly white blood cell, monocyte, and basophil counts. Shuffled controls showed negligible enrichment.

Together, these results establish that FOODIE footprints capture TF binding and enhancer– gene linkages more effectively than ATAC- and DNase-based approaches and are highly enriched for erythroid trait heritability. We therefore applied varTFBridge with K562 FOODIE footprints to 490,640 UK Biobank genomes to identify causal variants that alter TF-mediated gene regulation of erythroid traits.

### varTFBridge functionally prioritizes common noncoding variants by elucidating TF– mediated regulatory mechanisms

We performed GWAS for 13 erythroid traits across 490,640 UK Biobank participants of white European ancestry. After quality control, an average of 21,107,428 variants per trait (MAF ≥ 0.1%) were available. GWFM fine-mapping identified 251 credible variants (PIP ≥ 10%, PEP ≥ 70% for at least one erythroid trait) overlapping K562 FOODIE footprints (Fig. 3a).

**Fig. 3.**
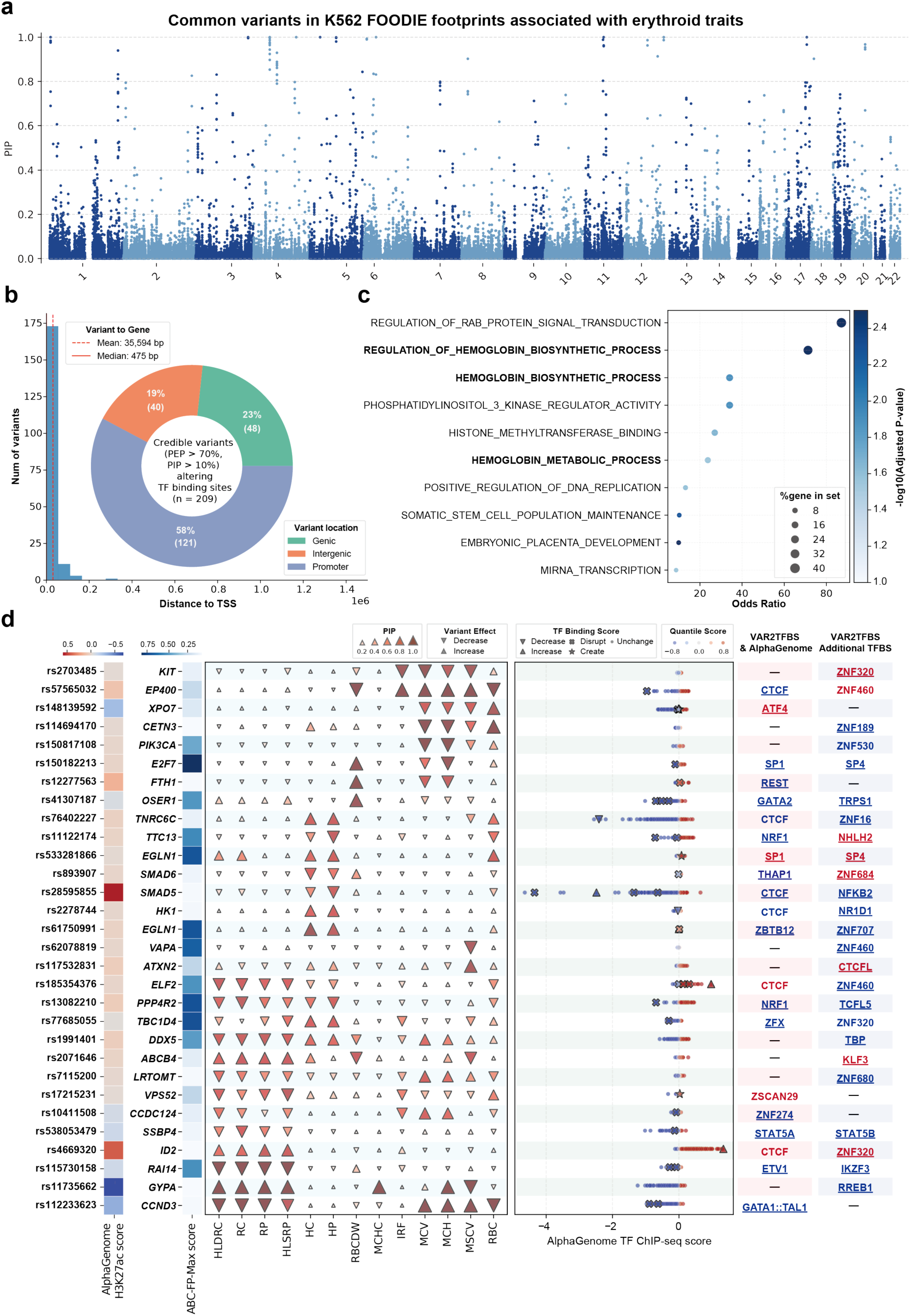
varTFBridge deciphers the impact of common variants on transcription-factor mediated function. **a,** Manhattan plots of fine-mapped common variants (MAF>0.1%) overlapping K562 FOODIE footprints across 13 erythroid traits. Posterior inclusion probabilities (PIPs) were estimated by Genome-wide fine-mapping using SBayesRC. Each point represents a footprint-overlapping variant plotted by genomic position. **b**, Pie chart depicting the proportion of credible set variants (PIP > 10% and PEP > 70%) by genomic annotation. “Genic” indicates variants within a gene body (intron); “Promoter” denotes variants within 500 bp of a gene’s transcription start site (TSS); “Intergenic” refers to regions without gene annotations. Gene annotations are based on ABC scores. Histogram showing the distribution of distances from variants to their predicted target genes, as determined by ABC-FP-Max. **c**, Gene Ontology (GO) enrichment of ABC-FP-Max predicted targets derived from erythroid trait–associated credible set variants (PIP > 10%). Each dot represents an enriched GO term. The x-axis indicates the odds ratio (OR) for enrichment; the y-axis lists the GO terms. Dot color signifies enrichment significance as −log10(adjusted P-value), and dot size corresponds to the percentage of genes in the GO term. **d**, Integrative demonstration of credible set variants (PIP > 70%) from trait association to TF-mediated regulatory function via VAR2TFBridge. The first column displays a heatmap indicating AlphaGenome-predicted variaant effects on K562 H3K27ac ChIP-seq (enhancer activity), with variant rsIDs on the y-axis. The second column shows a heatmap of ABC-FP-Max scores for variant-gene pairs, with the predicted target gene on the y-axis. The third column presents a heatmap of PIP values for each variant-trait pair, where triangle size denotes PIP magnitude and up/down triangles indicate positive/negative trait effects, respectively. The fifth column illustrates AlphaGenome-predicted variant variant effects on K562 TF ChIP-seq using scatter plots, where each point represents an AlphaGenome raw score for a specific TF, colored by its AlphaGenome quantile variant score. Up-triangles signify increased TF binding via VAR2TFBS, down-triangles decreased binding, crosses indicate motif disruption, and stars denote motif creation. The seventh column lists TFs whose binding sites are predicted to be altered by variants from VAR2TFBS with an AlphaGenome quantile score > 0.7. The last column identifies VAR2TFBS-predicted TFs whose binding sites are altered but are not in the AlphaGenome prediction list. TF names colored blue indicate decreased or disrupted binding (underlined for disruption), while red indicates increased or created binding (underlined for creation). If VAR2TFBS predicts multiple TFs for a variant, the one with the highest RNA-seq expression in K562 is listed; a hyphen indicates no TF was found by the specific strategy. A full list of TFs is in Supplementary Table 2, and erythroid trait names of aberration are detailed in the legend of Fig. 3.

We calculated an ABC-FP score for each footprint–gene pair within a 5 Mb window by multiplying estimated chromatin accessibility (ATAC-seq) by Hi-C contact frequency with the gene promoter (Methods). Pairs exceeding a threshold of 0.015 were defined as having footprint–gene linkage.

We applied VAR2TFBS to assess how credible set variants within TF footprints affect binding affinity. Using 879 vertebrate non-redundant PWMs from JASPAR 2024^30^, FIMO^29^ identified significant motif hits (p < 1 × 10⁻⁴) in both reference and alternative sequences. Changes in p-value between alleles were classified as TF binding disruption, creation, increase, or decrease (Methods).

VAR2TFBS identified 209 credible variants predicted to alter TF binding at ABC-FP footprints, of which 189 (90.4%) affect at least one expressed TF (K562 TPM > 0.5). These variants alter 1,154 predicted binding sites across 330 TFs. SP/KLF family sites account for 22% (n = 254), and CTCF/CTCFL sites for 3.7% (n = 43).

We assigned each variant to the gene with the highest ABC-FP score (ABC-FP-Max), yielding 209 variant–gene pairs involving 195 unique genes (Fig. 3b and Supplementary Table 1). Of these, 58% (n = 121) are in promoter regions and 19% (n = 40) in distal intergenic regions (median distance 475 bp; range 5 bp–3.33 Mb). GO enrichment analysis^40^ of target genes showed significant enrichment for hemoglobin biosynthesis and metabolism (Fig. 3c).

Individual variants affect a median of 4 TF binding sites, probably reflecting motif similarity across TF families. To quantify variant effects in the K562 context, we used AlphaGenome quantile scores, which rank the predicted allelic difference for each epigenomic feature (e.g., TF ChIP-seq, H3K27ac) against the distribution of effects from common variants genome-wide, thereby identifying variants with unusually large predicted effects (Methods). Combining VAR2TFBS and AlphaGenome, we annotated 28 TF binding sites (|quantile score| > 80%) altered by 30 high-confidence variants (PIP > 70%), including key erythroid regulators GATA1, GATA2, and TAL1^41^ (Fig. 3d and Supplementary Data 1). Because AlphaGenome covers only 269 TF ChIP-seq profiles, VAR2TFBS provides complementary predictions for TFs such as KLF5, KLF6, and KLF16 (Supplementary Table 2).

Having established the varTFBridge framework for common variants, we next asked whether it could be extended to rare noncoding variants, which have been largely inaccessible to regulatory interpretation.

### varTFBridge functionally prioritizes rare driver variants within erythroid trait– associated FOODIE footprints

We aggregated rare variants (MAF < 0.1%; carrier count > 30) within each K562 FOODIE footprint across 13 erythroid traits in 490,640 UK Biobank participants. Bonferroni-corrected burden testing identified 23 footprints significantly associated with at least one trait (Fig. 4a). Leave-one-variant-out analysis identified the driver variant within each significant footprint (Fig. 4b and Methods); as most associations were driven by a single variant (Supplementary Fig. 2), we designated that variant as the driver. In total, 18 rare driver variants were identified.

**Fig. 4.**
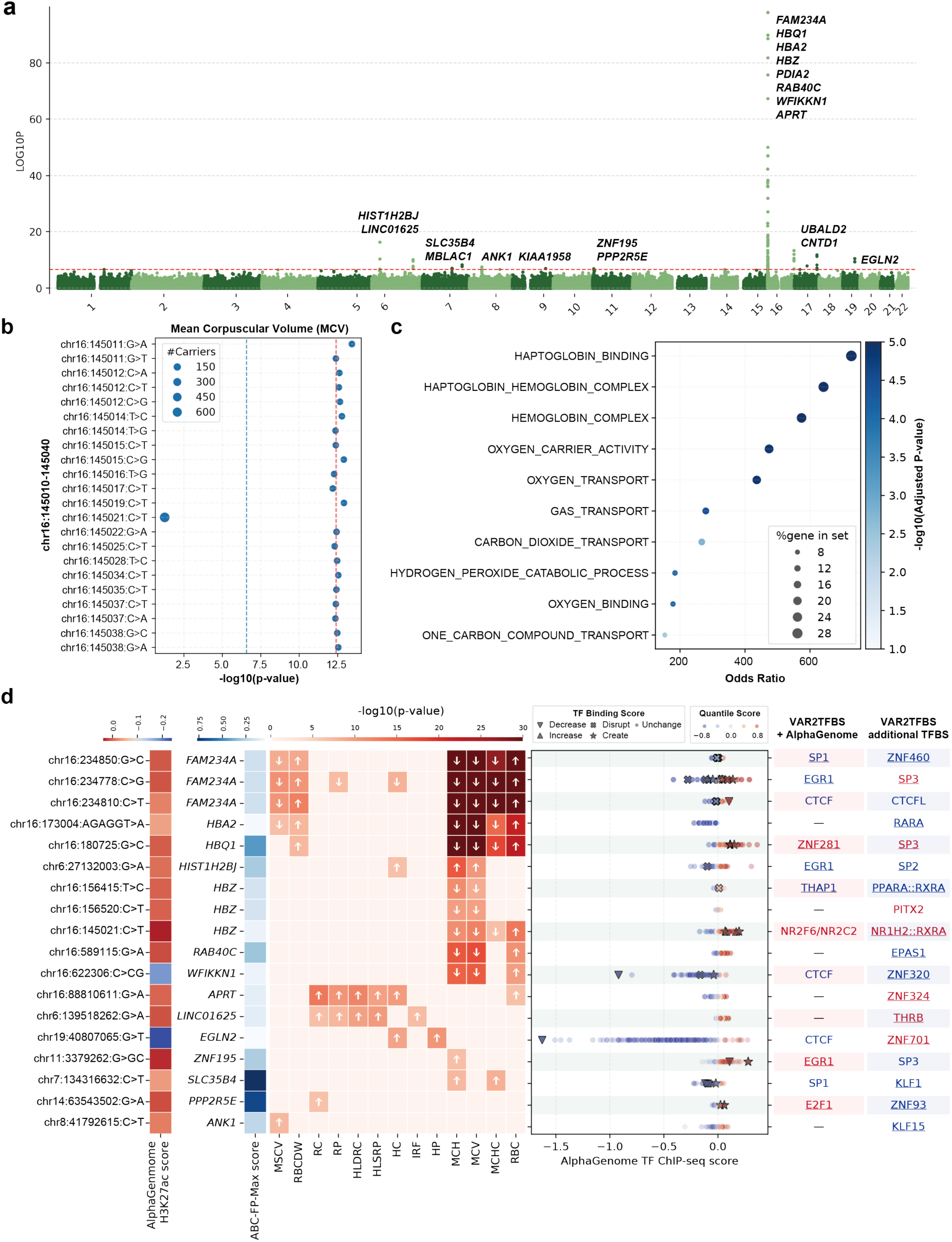
varTFBridge deciphers the impact of driver rare variants in erythroid-associated footprints on transcription-factor mediated function. **a.** Manhattan plots for FOODIE footprints-based burden test results for 13 erythroid traits, showing unadjusted two-sided P values derived from footprint-burden testing conducted in REGIENE and plotted on a -log10 scale. The x-axis denotes the genomic position of the center of the footprints. Scatters above the red dashed line are significant footprints (*P* < 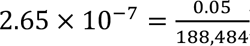). The ABC-FP-Max predicted target genes of the significant footprints are labeled. **b**, Scatter plot illustrating the leave-one-out analysis results on a significant footprint (chr16:145,010-145,040) associated with Mean Corpuscular Volume (MCV). Each scatter represents a rare variant (MAF < 0.1%) within this footprint, with the size of the scatter indicating the number of carriers of the variant. The blue dashed line indicates the significance threshold (6.57 = - log10(0.05/188,484)), and the red dashed line denotes the-log10(p-value) of footprint-based burden tests without variant masking. **c**, Gene Ontology (GO) enrichment of ABC-FP-Max predicted targets derived from significant footprints associated with erythroid traits. Each dot represents an enriched GO term, with the x-axis indicating the odds ratio (OR) for enrichment and the y-axis listing the GO terms. Dot color signifies enrichment significance as -log10(adjusted P-value), while dot size corresponds to the percentage of genes in the GO term. **d**, Integrative demonstration of top driver rare variants at significant FOODIE footprints, linking trait associations to transcription factor– mediated regulatory function. The first column heatmap indicates the AlphaGenome predicted effect on K562 H3K27ac ChIP-seq tracks, with the y-axis showing the genomic locus of the variants. The second column heatmap shows the ABC-FP-Max score of variant-gene pairs, with the y-axis showing the predicted target gene. The third column heatmap shows the-log10(p-value) of each variant-trait pair, where an up-arrow denotes a positive effect on the traits and a down-arrow denotes a negative effect. The fifth column shows AlphaGenome-predicted variant effects on TF ChIP-seq, where each scatter represents an AlphaGenome score of a specific TF, and the color indicates the AlphaGenome quantile variant score. Scatters marked with an up-triangle indicate the variant predicts to increase TF binding affinity via VAR2TFBS; a down-triangle indicates a decrease; a cross indicates disruption of the TF binding motif; and a star indicates creation of the TF binding motif. The seventh column lists the TF whose binding site is predicted to be altered by a variant using VAR2TFBS with an AlphaGenome quantile score > 0.85. The last column indicates the VAR2TBS identified TF whose binding site is altered by the variant but is not in the AlphaGenome prediction TF list. TF names colored in blue denote decreased or disrupted TF binding (underlined for disruption), and red denotes increased or created TF binding (underlined for creation). If VAR2TFBS predicted multiple TFs for a variant, the one with the highest RNA-seq expression in K562 is listed; a hyphen denotes no TF was found by the specific strategy. The full list of TFs is in Supplementary Table 3, and the full names of erythroid traits are detailed in the legend of Fig. 3.

ABC-FP-Max linked driver rare variants to 14 unique target genes, enriched for haptoglobin- and hemoglobin-related functions (Fig. 4c). VAR2TFBS and AlphaGenome annotated 35 TF binding alterations across 24 TFs (|quantile score| > 0.8; Fig. 4d and Supplementary Data 3), including erythroid regulators (ZBTB7A, ZNF148, NR2C2), genome-architecture factors (CTCF, SP1), and cell-fate TFs (EGR1, ETV6).

We then focused on the *HBZ*, a key component of primitive erythropoiesis that is highly expressed in K562 cells^42^. A previous study performed CRISPR perturbation to identify two *HBZ* enhancers^31^. We identified a driver rare variant (chr16:145021:C>T; Fig. 4b) residing in one of the enhancers (chr16:144,736-145,236), which is predicted by varTFBridge to regulate HBZ by altering the binding of NR2F6 (quantile score=0.981) and NR2C2 (quantile score = 0.944). NR2F6 (EAR-2) and NR2C2 (TR4) are orphan nuclear receptors with established roles in hematopoietic regulation: NR2F6 maintains clonogenic status in leukemia cells and its overexpression inhibits hematopoietic differentiation^43,44^, while NR2C2 directly represses embryonic β-type globin transcription and regulates erythroid cell proliferation and maturation^45,46^, making a predicted NR2C2 binding alteration at an HBZ enhancer biologically coherent.

### A high-confidence resource of variant–TF binding–gene regulatory hypotheses for erythroid traits

Having applied varTFBridge to both common and rare variants, we next assessed the combined output to define a high-confidence resource. Through progressive multi-layer filtering, varTFBridge distils the full spectrum of genetic variation into a prioritized set of variant–TF binding–gene regulatory hypotheses. For common variants, the pipeline begins with 13,065,104 GWFM SNPs across 15 blood traits and applies genome-wide fine-mapping to extract 60,450 credible set variants (PIP > 0.1) for 13 erythroid traits. Intersecting these variants with K562 FOODIE TF footprints retains 1,087 variants that fall within experimentally defined binding sites. VAR2TFBS then predicts 839 variants that alter the binding affinity of at least one TF, and ABC-FP-Max links each variant to its predicted target gene (ABC score > 0.015), yielding a final set of 209 common variants encompassing 1,154 variant–TF binding–gene regulatory hypotheses across 330 predicted TFs and 195 predicted target genes. For rare variants (MAF < 0.1%), footprint-based burden testing followed by leave-one-variant-out analysis identifies 23 driver variants within significant footprints; VAR2TFBS and ABC-FP-Max filtering produces 18 rare variants encompassing 141 regulatory hypotheses across 98 predicted TFs and 14 predicted target genes. The complete filtering funnel is summarized in Supplementary Data 4 and Supplementary Fig. 4. Detailed characterisation of the trait, TF, and gene distributions within this resource is provided in Supplementary Note 1.

To further prioritize the most reliable regulatory hypotheses from the full set of 227 variants produced by the pipeline described above, we applied three additional independent filters requiring convergent support: (1) TFBS altered: the variant must be predicted to alter TF binding affinity (Disrupt, Create, Increase, or Decrease); (2) TF expressed: the predicted TF must be expressed in K562 cells (RNA-seq TPM >= 1), confirming biological relevance in the cell type of interest; and (3) AlphaGenome epigenomic concordance: at least one of 12 K562 epigenomic markers (10 histone modifications, ATAC-seq, and DNase-seq) must show a strong AI-predicted variant effect (|quantile score| > 0.95), providing independent computational evidence for a regulatory impact at the variant position.

This three-way intersection yielded 113 high-confidence variants (104 common, 9 rare) targeting 108 unique predicted genes and involving 64 unique predicted TFs (Supplementary Data 5). Among high-confidence common variants, predicted TF binding disruption (57) and de novo creation (44) were the dominant alteration types, with key erythropoiesis regulators including CTCF (17 variants, the most frequently altered TF), KLF1 (7 variants), SP1, SP3, CTCFL, and EGR1 (4 variants each). Of the 18 rare variants, 9 passed all three filters, including driver variants at the alpha-globin locus (HBA2, HBQ1, HBZ on chr16) and variants targeting SLC35B4 (predicted KLF1 disruption) and HIST1H2BJ (predicted EGR1 disruption).

The complete resource of 113 variants, with all evidence columns and high-confidence status (Supplementary Data 5). All variant–TF binding–gene predictions remain computational hypotheses that require experimental validation; the convergent-evidence filter is designed to prioritize the most promising candidates for targeted functional studies such as CRISPR perturbation, luciferase reporter assays, or allele-specific binding experiments. The three filters rely on generalizable pipeline components (VAR2TFBS, RNA-seq expression, and AlphaGenome), ensuring that the framework can be directly applied to other cell types as FOODIE data become available.

### varTFBridge predicts the molecular mechanism underlying GATA1/TAL1-dependent CCND3 enhancer regulation

Among the 113 high-confidence variants, rs112233623 stands out as a uniquely informative test case: it carries PIP = 1.0 across 10 erythroid traits, resides within a locus where prior functional studies established the variant-to-gene link but left the causal TF mechanism unresolved, and thus provides an opportunity to assess whether varTFBridge can independently recover and extend known biology. To demonstrate the mechanistic resolution enabled by varTFBridge, we examined the well-characterized *CCND3* locus on chromosome 6p21.1, where prior studies established the variant-to-gene link and functional significance but left the molecular mechanism unresolved. CCND3 encodes cyclin D3, which regulates the number of cell divisions that erythroid precursors undergo during terminal differentiation, thereby controlling both erythrocyte size and number. A previous study identified an erythroid-specific enhancer within *CCND3* intron 3 and confirmed its activity via luciferase reporter assays in K562, with no activity in non-erythroid cells (293, HeLa, MCF-7)^47^. ChIP-seq in K562 and primary erythroid progenitors demonstrated GATA1 and TAL1 binding at this enhancer, yet no consensus TF binding motif was identified at the sentinel variant rs9349205. Subsequent fine-mapping resolved a second causal variant, rs112233623 (PP = 0.99), located 161 bp from rs9349205 (PP = 0.94) within the same enhancer^48^. Luciferase assays in K562 confirmed that each variant independently affects enhancer activity, and ATAC-seq in primary hematopoietic progenitors showed both variants reside within erythroid-specific accessible chromatin. Despite these functional validations, the specific TF binding motif disrupted, and the downstream regulatory cascade remained unknown.

varTFBridge proposes a specific molecular mechanism for this locus, supported by multiple independent lines of evidence. Using K562 FOODIE footprints and motif scanning (VAR2TFBS), our analysis predicts that rs112233623, not the sentinel rs9349205, disrupts a GATA1/TAL1 heterodimer binding motif (JASPAR MA0140.2). This prediction is corroborated by K562 ENCODE ChIP-seq data: GATA, TAL1, and the histone acetyltransferase EP300 all show binding peaks directly overlapping rs112233623 (ENCODE encRegTfbsClusteredWithCells), confirming that these factors physically occupy the variant position. The motif prediction thus explains the paradox from previous study^47^: the GATA1 and TAL1 ChIP-seq peaks are real and centered on the enhancer, but the cognate co-binding motif lies at rs112233623, 161 bp from the sentinel and well within the spatial resolution of a single ChIP-seq peak. The variant is associated with 10 erythroid traits (PIP = 1.0 across RBC, MCV, MCH, MSCV, RBCDW, RC, RP, HLDRC, HLSRP, MCHC) (Fig. 5a). AlphaGenome variant effect predictions independently corroborate this mechanism, predicting concordant loss of GATA1, TAL1, and EP300 binding alongside reduced H3K27ac and chromatin accessibility at the variant position (Fig. 5b, Supplementary Note 2 and Supplementary Data 2).

**Fig. 5.**
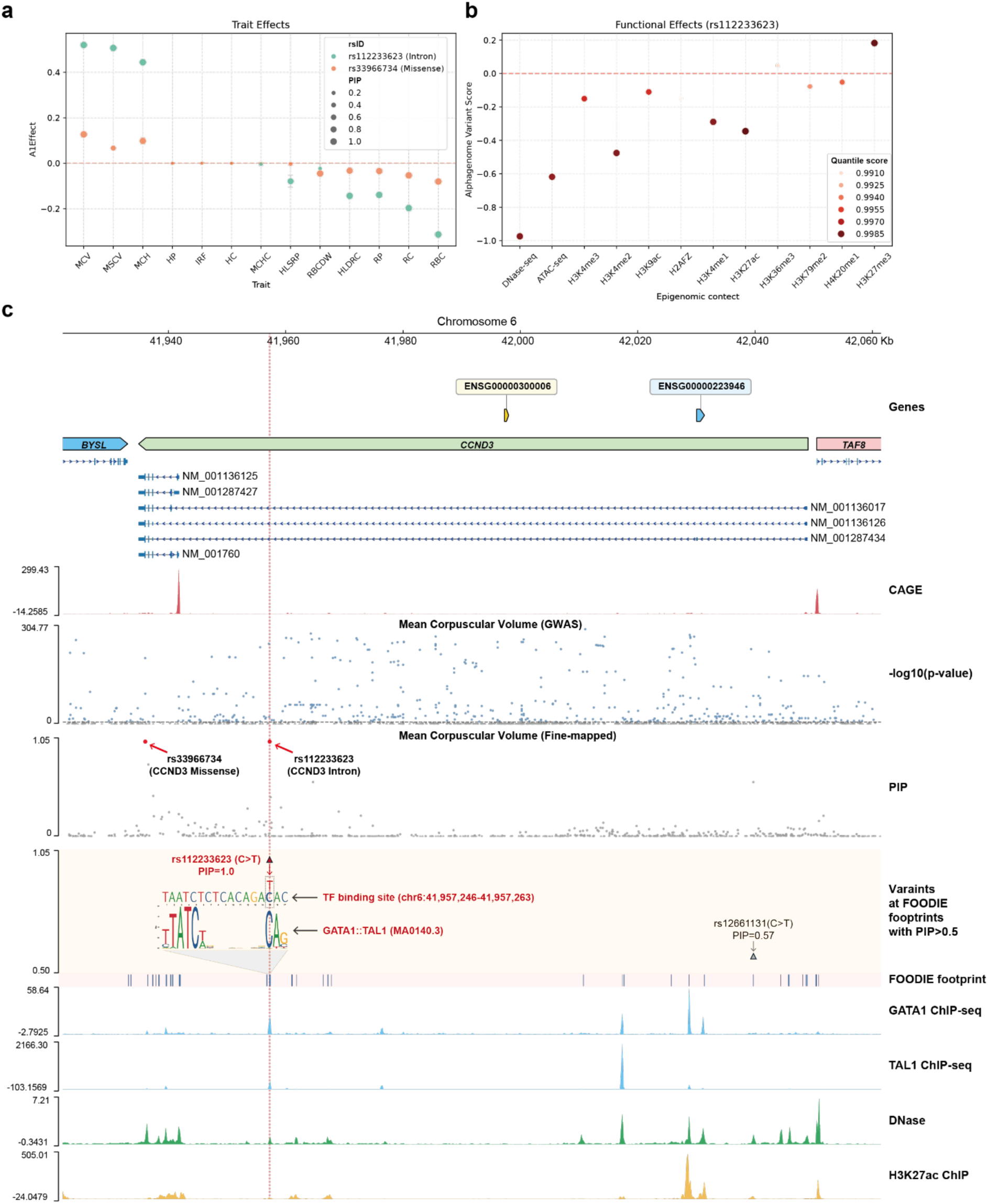
varTFBridge identifies rs112233623 as a causal regulatory variant predicted to modulate CCND3 expression by disrupting a GATA1/TAL1 binding site, thereby underlying erythroid phenotypic variation. **a,** Scatter plot showing the estimated effect size (BETA) and Posterior Inclusion Probability (PIP) of fine-mapped variants, rs112233623 and rs33966734, across 13 erythroid traits. A positive effect size indicates the variant increases the trait, while a negative effect decreases it. The size of each scatter point denotes its PIP value, and color differentiates between the variants. Error bars represent one standard error of the effect size, and the red dashed line indicates zero effect on the trait. **b,** Scatter plot displaying the AlphaGenome variant score of rs112233623 based on 11 histone marker ChIP-seq and 2 chromatin accessibility tracks. The color of each scatter indicates the absolute AlphaGenome quantile score, reflecting the variant’s effect compared to background common variants. The red dashed line denotes a zero score from AlphaGenome, signifying no effect on the specific epigenome profiles. **d**, Genome browser view illustrating Mean Corpuscular Volume (MCV) associated variants at the *CCND3* genomic locus (chr6:41,922,000-42,061,500) with multiple epigenomic tracks. The first row displays the *CCND3* gene along with its different isoforms and two nearby genes. The second row presents GWAS signals, where each scatter represents a variant (MAF > 0.1% from 490,640 UK Biobank genomes) with-log10(p-value) from GWAS on the y-axis. Blue scatters denote significant variants (p-value > 5e-8, indicated by the grey dashed line). The third row shows fine-mapped signals for the same variants, with PIP on the y-axis. Red scatters represent the two credible set variants with PIP = 100%, namely rs112233623 (located in a distal enhancer upstream of CCND3) and rs33966734 (located in introns). The fourth row demonstrates variants at K562 FOODIE footprints (PIP > 50%) and specifically illustrates how rs112233623 (C-to-T substitution) disrupts the GATA1/TAL1 binding site. The sequence shown is extracted from the hg38 reference genome (chr6: 41,957,246-41,957,263), and the GATA1::TAL1 (JASPAR ID: MA0140.3) motif logo is sourced from the JASPAR 2025 database. The fifth row displays all K562 FOODIE footprints within this genomic region. The subsequent four rows present genomic tracks of GATA1 ChIP-seq, TAL1 ChIP-seq, DNase, and H3K27ac ChIP-seq signals from ENCODE, in order. The red vertical dashed line denotes the TSS of *CCND3*.

The convergence of four independent lines of evidence: (i) motif-based prediction of GATA1/TAL1 disruption, (ii) ENCODE ChIP-seq confirming GATA1, TAL1, and EP300 occupancy at the variant position, (iii) AlphaGenome predicting concordant loss of these factors and downstream chromatin marks, and (iv) prior luciferase assays confirming that this variant alters enhancer activity^48^, supports a coherent mechanistic model: rs112233623 disrupts the GATA1/TAL1 co-binding motif at an erythroid-specific enhancer, leading to loss of EP300 co-activator recruitment, erasure of the H3K27ac active enhancer mark, chromatin closing, and ultimately reduced *CCND3* expression. This connects the variant to the cell-biological phenotype of fewer terminal erythroid divisions, decreased RBC count, and reciprocally increased erythrocyte size. While prior work established that variants at this locus affect enhancer activity and linked them to *CCND3*, varTFBridge provides the first prediction of the specific TF motif disrupted and proposes a complete regulatory cascade from motif disruption to chromatin remodeling, demonstrating the framework’s capacity to generate reliable, mechanistically detailed hypotheses by integrating high-resolution TF footprinting, motif-based binding predictions, ChIP-seq occupancy data, and AI-based variant effect scoring. (Fig. 5c and Supplementary Fig. 5)

## Discussion

Despite nearly two decades of GWAS, functionally characterizing noncoding variants and mapping them to target genes remain a major bottleneck. This challenge is compounded for rare variants, which require higher-resolution functional annotations than coding-region burden tests. Here, we show that FOODIE footprints provide the necessary resolution: they improve TF binding measurement and target gene prediction over ATAC-seq and DNase-seq and yield substantially higher heritability enrichment for erythroid traits in K562 cells, with raw footprints covering 0.12% of the genome yet capturing up to 103-fold enrichment. Building on these properties, varTFBridge integrates FOODIE footprinting with motif scanning, enhancer–gene linkage, and deep-learning variant effect prediction to map noncoding variants to TF-mediated regulatory mechanisms across the allele frequency spectrum.

Applied to 490,640 UK Biobank genomes, varTFBridge produced a regulatory resource of 227 variant–TF binding–gene linkages for 13 erythroid traits, from which convergent-evidence filtering prioritized 113 high-confidence candidates. The framework independently recapitulated the causal variant rs112233623 and resolved its mechanism as disruption of GATA1/TAL1 co-binding at a CCND3 enhancer, demonstrating that varTFBridge can both recover known biology and generate testable regulatory hypotheses.

varTFBridge is, to our knowledge, the first framework to systematically investigate both common and rare noncoding variants through TF footprinting. The combination of VAR2TFBS, ABC-FP-Max, and AlphaGenome provides synergistic advantages. While AlphaGenome cannot predict the effects of distal variants on target genes, ABC-FP-Max provides robust variant-to-gene links up to 5 Mb away. Conversely, AlphaGenome’s TF binding predictions are limited to 128 bp resolution due to ChIP-seq constraints, whereas VAR2TFBS achieves base-resolution predictions through FIMO motif scanning within FOODIE footprints. When these independent approaches converge on the same prediction, confidence in the regulatory hypothesis increases substantially, making it a high-priority target for experimental follow-up.

Our study has several limitations that highlight areas for future work. First, varTFBridge only considers variants residing within TF binding sites, thereby missing variants flanking the core motif that might also affect TF binding affinity. Second, some co-regulators do not bind directly to the DNA sequence; our model remains limited in predicting this type of co-regulation, though a recent model, Chromnitron^49^, might help bridge this gap. Third, varTFBridge assumes a single causal gene per variant, although enhancers containing disease variants often have highly pleiotropic effects. Fourth, AlphaGenome can only predict a limited number of TFs, which hinders the extension of our framework to other tissues and cell lines. Integrating ChromBPNet^50^ into our framework could potentially address this limitation, as it can be easily trained on FOODIE data from new cell lines and tissues to predict variant effects. Lastly, while our multi-evidence approach substantially increases the credibility of predictions, the variant–TF binding–gene hypotheses generated by varTFBridge remain computational predictions that require experimental validation. We provide this resource as a prioritized starting point, not a substitute, for targeted functional studies.

In summary, our approach paves the way for a comprehensive mapping of noncoding variants across the allele frequency spectrum onto TF-mediated gene regulatory networks. While this study utilizes experimental data from K562 cells to model erythroid traits underlying anemia and related disorders, we anticipate that varTFBridge will be broadly applicable to other cell lines and tissues as FOODIE footprinting data becomes available, enabling the community to systematically decode the regulatory grammar of noncoding disease variants through targeted experimental validation.

## Methods

### Ethics

The UK Biobank study has ethical approval from the North West Multi-centre Research Ethics Committee (REC reference: 13/NW/0157) as a Research Tissue Bank (RTB). This approval is renewed every five years, with successful renewals in 2016 and 2021. All participants provided informed consent, and the present analyses were conducted under UK Biobank application number 477126.

### Cell Preparation

Human leukemia cell line K562 and human lymphoblastoid cell line GM12878 were grown with RPMI 1640 medium (cat. no. 11875093, Gibco) containing 10% FBS (cat. no. 10099141C, Gibco) in a 37 °C incubator supplied with 5% CO2. Cells were rinsed once with 1 mL of DPBS (cat. no. 14040117, ThermoFisher) and resuspended in 1 mL DPBS before use.

### FOODIE library preparation

The FOODIE libraries were constructed using the commercial Hyperactive FOODIE Library Prep Kit (cat. no. TD721, Vazyme) following the manufacturer’s instruction, which is similar to our previously reported protocol^22^. Briefly, approximately 50,000 cells were counted, washed with TW Buffer, and centrifuged. The cell pellet was resuspended in pre-chilled Lysis Buffer to facilitate permeabilization. Following permeabilization, nuclei were collected via centrifugation and resuspended in Tagmentation Mix for the Tn5 transposition reaction. The reaction was incubated at 37 °C for 30 min on a thermomixer with shaking at 800 rpm. Subsequently, the tagmentation reaction was terminated by adding pre-chilled Wash Buffer, followed by centrifugation. The isolated nuclei were then resuspended in a Deamination Mix supplemented with TruePrep DddB Enzyme (cat. no. S802-01, Vazyme) and incubated at 37 °C for 10 min. The reaction was halted by the addition of Stop Buffer. Finally, DNA was purified using the provided magnetic beads, followed by PCR amplification using the kit-supplied DNA polymerase and primers.

### FOODIE data processing and footprint calling

We followed the procedure described in the original FOODIE study^22^. Briefly, raw reads were trimmed with Trim Galore (v0.6.10)^51^, and aligned to the human reference genome hg38 and deduplicated with Bismark (v0.24.2)^52^. Customized scripts were then applied to perform deamination/conversion, followed by calculation and normalization of the conversion ratio at each genomic position. The adjusted conversion ratios together with the corresponding sequencing depths were used for open region identification and transcription factor footprint calling. Footprints with P values < 1 × 10⁻⁷ were retained for downstream analyses. The scripts are available at https://github.com/sunneyxie-lab/bulk-foodie-pipeline.

### ATAC-seq and DNase-seq footprint calling and processing

We leveraged HINT^33^ (v1.0.2), a Hidden Markov Models–based method, to derive transcription factor footprints from ATAC-seq data in K562 cells. We downloaded the K562 ATAC-seq BAM file (ENCFF077FBI) and its corresponding narrow peak file (ENCFF333TAT) from ENCODE and processed them with HINT’s default settings. K562 DNase footprints (ENCFF771QAQ) in BED format were also obtained from ENCODE.

### Genome-wide association testing of erythroid traits in UK Biobank

We applied the same GWAS framework as described in our previous analysis^53^ for UK Biobank. In addition to UK Biobank’s standard central quality control, we restricted analyses to participants of white European genetic ancestry. This was determined using k-means clustering of the first four principal components derived from genome-wide SNP genotypes. Individuals assigned to the white European cluster, but who self-reported a non-European ancestry were excluded. To avoid confounding due to relatedness, only unrelated individuals were included in the association analyses. After all quality control filters, a total of 490,640 participants with both genotype and phenotype data were eligible for inclusion. We further removed variants with minor allele frequency (MAF) < 0.1%. Genome-wide association analysis was performed using REGENIE v3.3^54^, implemented within the UK Biobank Research Analysis Platform and following recommendations for UK Biobank analyses (https://rgcgithub.github.io/regenie/recommendations/). The association model analyzed 13 erythroid-related traits as outcome variables: hematocrit (HP), hemoglobin concentration (HC), high light scatter reticulocyte count (HLSR), high light scatter reticulocyte percentage (HLSRP), immature reticulocyte fraction (IRF), mean corpuscular hemoglobin (MCH), mean corpuscular hemoglobin concentration (MCHC), mean corpuscular volume (MCV), mean sphered corpuscular volume (MSCV), red blood cell count (RBC), red cell distribution width (RBCDW), reticulocyte count (RC), and reticulocyte fraction of red cells (RP). Covariates included genotyping array, sex, age at questionnaire completion, and the first ten principal components of genetic ancestry.

### Genome-wide fine-mapping for identifying credible set variants

Credible set variants were derived from genome-wide fine-mapping (GWFM) with the SBayesRC^55^ framework, which implements a Bayesian hierarchical mixture model that integrates genome-wide linkage disequilibrium (LD) patterns and functional genomic annotations to estimate posterior inclusion probabilities (PIPs) for SNPs. Genome-wide LD was captured using precomputed eigen-decomposition reference panels provided with GCTB^56^, ensuring consistency with the assumed genetic architecture. Functional annotations were incorporated from Gazal et al.^39^, encompassing 96 genomic features relevant to complex trait architecture, including coding and conserved regions. Fine-mapping was performed genome-wide. Local credible sets (LCS) were constructed for each candidate causal variant by aggregating variants in LD (*r*^2^ > 0.5) until their cumulative PIPs reached 70%, ensuring the LCS explains more heritability than a random SNP set of the same size. In this study, variants with PIP ≥ 10% and residing in an LCS with cumulative PIP ≥ 70% were considered likely causal, following previous studies^8,27^.

GWFM (v2.5.4) was downloaded from https://gctbhub.cloud.edu.au/software/gctb/#Download. Summary statistics from 13 erythroid trait GWAS were used to construct credible sets. Variants were mapped to reference SNP identifiers (rsIDs) using the dbSNP release^57^ (build b157) based on the GRCh38 human genome assembly (GCF_000001405.40). To align with the GWFM database version, genomic coordinates were lifted over from hg38 to hg19 using LiftOver. The processed hg19 summary statistics were then converted to the required .ma format, and the --impute-summary option was employed for quality control and imputation of summary statistics for SNPs present in the LD reference but not in the GWAS data. Finally, GWFM analyses were run with the --gwfm RC option, implementing the SBayesRC model.

### FOODIE footprint-based test and leave-one-variant-out analysis for rare variants

To identify rare germline variants (MAF < 0.1%) on FOODIE footprints associated with erythrocyte traits, we performed footprint-based burden tests across chromosomes 1–22 in 490,640 UK Biobank (UKBB) participants with whole-genome sequencing data. Briefly, 490,640 participants were sequenced at an average depth of 32.5× on the Illumina NovaSeq 6000. The WGS generation and processing pipeline for UKBB participants is described in the recent study^21^. We annotated the variants with “FILTER=PASS” from ML-Corrected DRAGEN whole genome sequencing (WGS) in PLINK2 format using the ENSEMBL Variant Effect Predictor (VEP) v113.0, and analyses were restricted to rare qualifying variants (MAF < 0.1%). Footprints with <30 qualifying variant carriers were excluded from each burden mask. Accordingly, the footprint-wide significance threshold was set using Bonferroni correction at: 0.05/188,484 = 2.65 × 10^−7^.

### Stratified linkage disequilibrium score regression

To identify blood cell traits with heritability enrichment in K562 FOODIE footprints, we employed stratified linkage disequilibrium score regression (S-LDSC)^58^. All GWAS data were preprocessed using the ‘munge_sumstats.py’ script. Variants in the Major Histocompatibility Complex (MHC) region were excluded from the analysis. In addition to K562 FOODIE footprints, ATAC-seq (ENCFF333TAT) and DNase-seq narrow peaks and footprints (ENCFF274YGF, ENCFF771QAQ) were downloaded from ENCODE^32^. All TF footprint annotations were extended by 50 bp on each side to ensure sufficient SNP coverage for robust S-LDSC estimation; ATAC-seq peaks were used without extension. Linkage disequilibrium (LD) scores were calculated for each annotation using the 1000 G Phase 3 European population^59^. The heritability enrichment of each annotation for a given trait was computed by adding the annotation to the baseline LD score model (v2.0) and regression for HapMap3 SNPs. These analyses used v1.0.1 of the LDSC package.

### ABC-FP-Max predictions for linking variants to the target gene

To link variants to their target genes using ABC-FP-Max, we first adapted the existing ABC model (accessible at https://github.com/broadinstitute/ABC-Enhancer-Gene-Prediction) to predict footprint–gene connections within specific cell types, thereby constructing ABC-FP models. The original ABC model was designed to link chromatin accessibility elements to genes, initially by calling peaks in chromatin accessibility data using MACS2. In contrast, for connecting FOODIE footprints to genes, we replaced the peak-calling step with FOODIE footprint calling. Following footprint identification, we extended each footprint by 50-bp on either side and merged them into a single element, recognizing that closely located footprints might co-regulate the same genes and larger windows can better count reads in ATAC-seq and H3K27ac ChIP-seq. Subsequently, we followed the original ABC models to estimate chromatin accessibility from ATAC-seq data on these footprint-based elements and to assess chromatin conformation (Hi-C) between these elements and the promoters of potential target genes, as previously described. Specifically, the activity of each footprint-based element is calculated based on ATAC-seq experiments and then quantile-normalized to the distribution observed in K562 cells. The footprint-gene contact is determined using cell-type specific Hi-C data from 4DN (4D Nucleome Data Portal). Finally, we compute the ABC-Max score for each footprint-gene pair as the product of activity and contact, normalized by the product of activity and contact for all elements within 5 Mb of that gene.

To calculate the ABC-FP-Max score for variant-gene pairs, we took the predictions from the ABC-FP model and applied the following processing steps. First, we considered all distal footprint–gene connections with an ABC-FP score ≥ 0.015 and all distal or proximal promoter–gene connections with an ABC score ≥ 0.1 (following the ABC-Max model setting). Second, we overlapped the variants with the ABC-FP nominated footprints. For each variant, the target gene was assigned as the one with the highest ABC-FP score among all target genes in the list; this top ABC-FP score is designated as the ABC-FP-Max score, quantifying the predicted variant-gene pair.

### Assessing fold excess overlap with direct measurements of TF ChIP-seq

We downloaded TF ChIP-seq data from ENCODE (https://hgdownload.soe.ucsc.edu/goldenPath/hg38/encRegTfbsClustered/encRegTfbsClusteredWithCells.hg38.bed.gz). We selected peaks annotated with K562 cell line and created a bed file with the union of ChIP-seq peaks from all TFs assayed. We then calculated the excess overlap between an annotation (A) and the ChIP-seq peaks (B) as

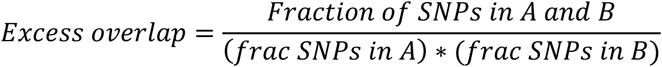

We calculated SDs using a block-jackknife, dividing the genome into 200 blocks of equal genomic size^14^.

### VAR2TFBS for predicting variant effects on TF binding sites

Quantifying the impact of genetic variants within FOODIE footprints necessitates assessing transcription factor (TF) binding affinity. VAR2TFBS employs a scoring procedure based on FIMO (https://github.com/jmschrei/memesuite-lite/blob/main/memelite/fimo.py), which identifies statistically significant hits by scanning Position Weight Matrix (PWM) of a certain TF against one-hot encoded DNA sequences. This process involves convolving PWMs across DNA sequences and converting the resulting scores into p-values using an exact background distribution derived from the PWM. In this study, we collected the PWMs from JASPAR 2024 core non-redundant vertebrates (https://jaspar.elixir.no/download/data/2024/CORE/JASPAR2024_CORE_vertebrates_non-redundant_pfms_meme.txt). To assess transcription factor (TF) binding within a FOODIE footprint, the sequence is first extended by 30 bp on each side. This extended reference sequence is then subjected to FIMO scanning, with TF binding sites identified by considering all Position Weight Matrices (PWMs) exhibiting a motif p-value less than 1 × 10^−4^.

Subsequently, each genetic variant is intersected with these identified TF binding sites, and FIMO scanning is performed on sequences incorporating the alternate allele. The resulting changes in TF binding are categorized into four types: creation, denoting a *de novo* TFBS (p-value < 1 × 10^−4^) in the alternate sequence; increase, where an existing TFBS shows a decreased (more significant) p-value in the alternate sequence; disruption, defined as an existing TFBS in the reference sequence having a p-value ≥ 1 × 10^−4^ in the alternate sequence; and decrease, signifying an existing TFBS with an increased (less significant) p-value in the alternate sequence.

In general, VAR2TFBS takes as input a reference genome, a variant list in BED file format, FOODIE footprints also in BED file format, and a collection of PWMs. It produces as output a comprehensive list of variants that overlap with FOODIE footprints, detailing their respective effects on transcription factor binding sites.

### AlphaGenome for variant effect scoring

We leveraged AlphaGenome (v0.5.1) to predict variant effects on cell-type–specific epigenomic profiles. The approach generates modality-specific predictions for reference (REF) and alternate (ALT) sequences, which are then compared to assess the regulatory impact of each variant across modalities. The model analyzes sequences up to 1 million bases and delivers single-base-pair resolution for ATAC-seq, DNase-seq, and TF ChIP-seq, and 128-bp resolution for histone modification ChIP-seq and TF binding. Variant scoring is performed by generating REF and ALT predictions within a modality-specific window around the variant: a 502-bp window centered on the variant for ATAC-seq, DNase-seq, and TF ChIP-seq, and a 2001-bp window for histone modification ChIP-seq. Predictions are aggregated across the window using log2-ratio of summed signals to yield a raw variant score: 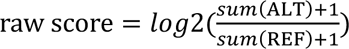.

Raw variant scores were normalized to empirical quantiles derived from common variants (MAF > 1% in GnomAD v3) to enable cross-track comparisons. For scores with directional effects, quantile probabilities were linearly transformed to the range [−1, 1], where the 0th, 50th, and 100th percentiles map to −1, 0, and +1, respectively. We report both raw and quantile scores across two chromatin accessibility modalities (ATAC-seq and DNase-seq) and 269 histone modification tracks in K562 cells.

### Convergent-evidence filtering for high-confidence variant prioritization

To prioritize the most reliable regulatory hypotheses, we applied three independent convergent-evidence filters to the full set of 227 pipeline variants. First, each variant must be predicted by VAR2TFBS to alter TF binding affinity (classified as Disrupt, Create, Increase, or Decrease). Second, the predicted TF must be expressed in K562 cells (RNA-seq TPM ≥ 1; ENCODE accession ENCFF485RIA), confirming biological relevance in the cell type of interest. Third, at least one of 12 K562 epigenomic tracks scored by AlphaGenome (10 histone modifications, ATAC-seq, and DNase-seq) must show a strong predicted variant effect (|quantile| > 0.95), providing independent computational evidence for regulatory impact. The intersection of all three filters yielded 113 high-confidence variants (104 common, 9 rare) targeting 108 genes through 64 TFs.

## Data availability

The UK Biobank phenotype and genome data described here are publicly available to registered researchers through the UK Biobank data access protocol. The FOODIE data is available at https://github.com/JasonLinjc/varTFBridge/tree/main/data.

## Code availability

The code used to develop varTFBridge, perform the analyses and generate results in this study is publicly available and has been deposited in https://github.com/JasonLinjc/varTFBridge under the MIT License.

## Acknowledgments

We thank the participants and investigators in the UKBB study (resource application nos. 477126) who made this work possible.

## Author contribution

This project is supported by Changping Laboratory. Y.Z. and X.X. conceived and supervised the study. J.L. developed varTFBridge and led downstream computational analyses, including model benchmarking and case studies. W.D. performed FOODIE experiments; C.X. conducted FOODIE footprint calling. J.L. and J. Zhang performed genome-wide fine-mapping. J.L., J. Zhao, and K.M. conducted GWAS and footprint-based burden tests. J.L., E.H., and J. Zhang performed S-LDSC analyses. Y.Z., J.L., X.X., C.X. and W.D. drafted the manuscript, with input from all authors.

