## Supplementary Figures for "Genome-wide maps of transcription factor footprints identify noncoding variants rewiring gene regulatory networks"

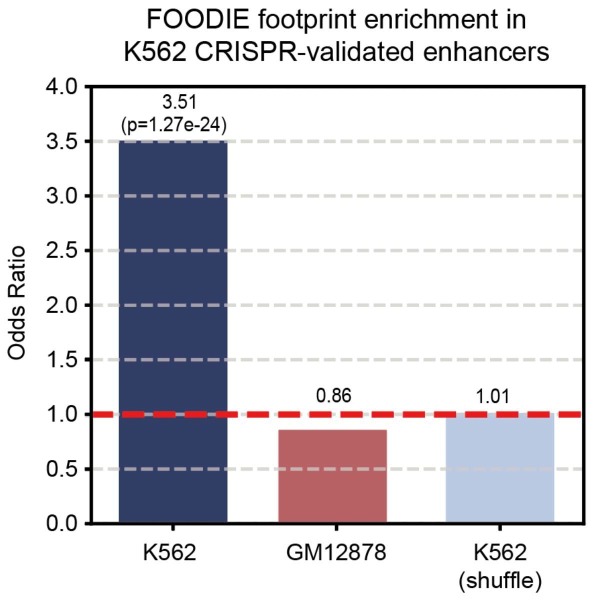


**Supplementary Figure 1.** Enrichment of K562 FOODIE footprints in CRISPR-validated enhancers. Bars show the fraction of CRISPR-validated enhancer regions (CRISPR Benchmark) overlapping K562 FOODIE footprints, GM12878 FOODIE footprints, and randomly shuffled K562 footprints.

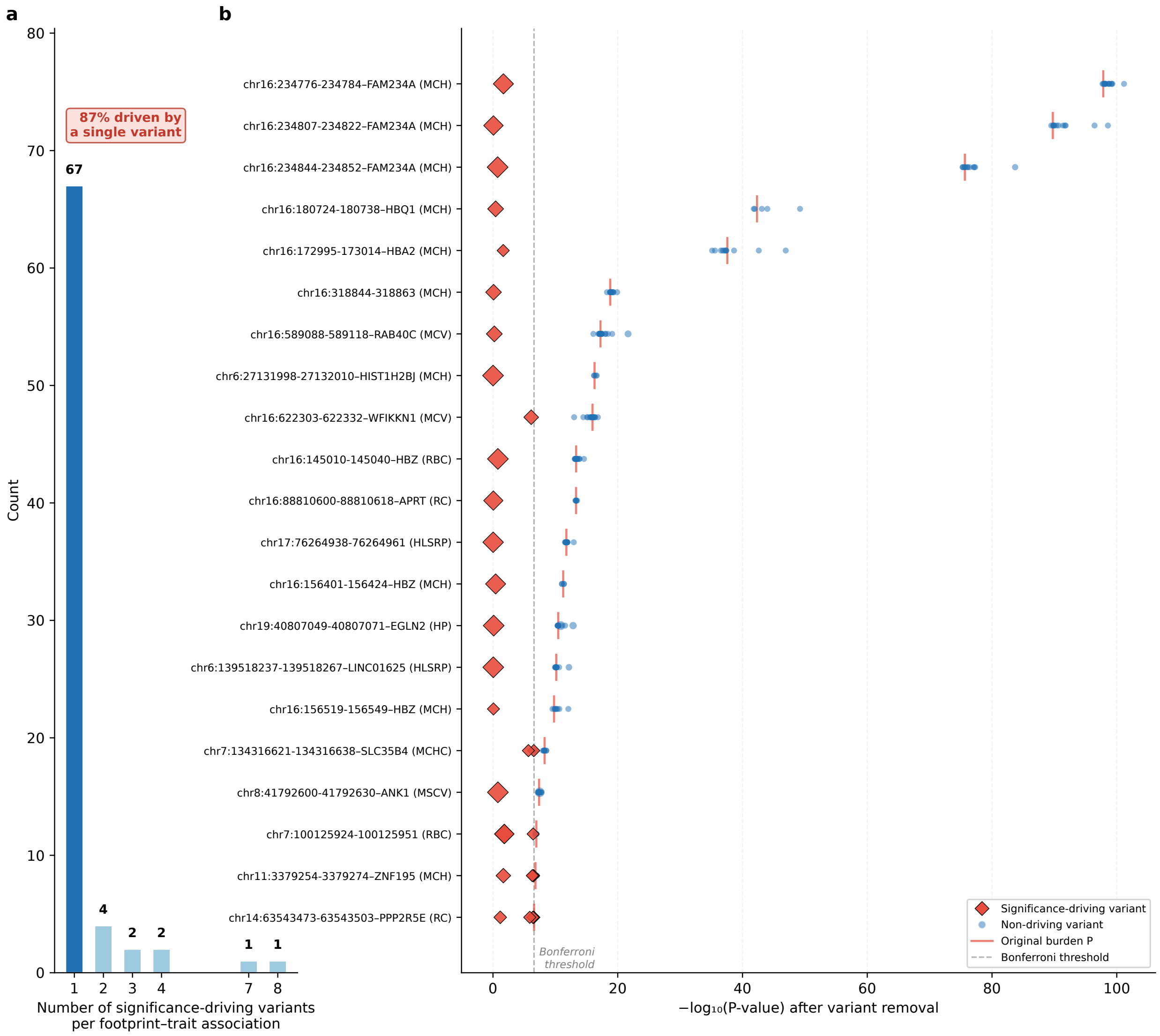


**Supplementary Figure 2.** Most erythroid trait–associated FOODIE footprints are driven by a single rare variant. a, Bar plot showing the distribution of the number of significance-driving variants per significant footprint–trait association (n = 77 associations across 21 unique footprints). A significance-driving variant is defined as one whose removal from the footprint-based burden test causes the footprint to lose statistical significance ($P > 2.65 \times 10^{-7}$). 87% (67/77) of associations are driven by a single variant. b, Leave-one-variant-out analysis results for 21 significant footprints, each shown for its most significant trait. Each point represents a rare variant within the footprint; the x-axis shows the $-\log_{10}$(P-value) of the burden test after removing that variant. Red diamonds denote significance-driving variants whose removal causes loss of significance; blue circles denote non-driving variants. Red vertical bars indicate the original burden test P-value (all variants included). The grey dashed line marks the Bonferroni-corrected significance threshold ($P = 0.05/188,484 = 2.65 \times 10^{-7}$). Y-axis labels indicate the footprint genomic coordinates, ABC-FP-Max predicted target gene, and the most significantly associated erythroid trait in parentheses. Full trait names are detailed in the legend of Fig. 2.

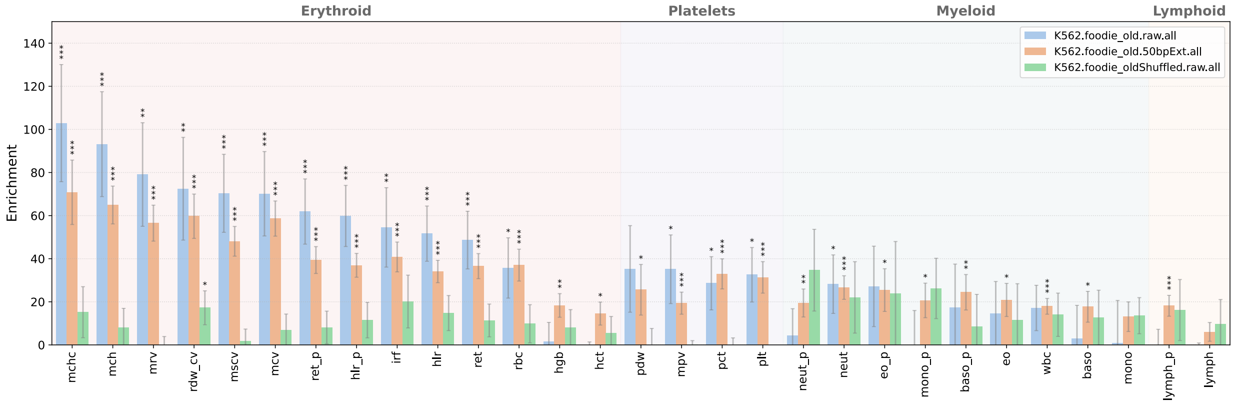


**Supplementary Figure 3.** S-LDSC heritability enrichment of raw and 50 bp-extended FOODIE footprints in K562. Partitioned S-LDSC enrichment estimates (fold enrichment over genome-wide baseline) are shown for 13 erythroid and 2 lymphoid traits. Extended footprints (50 bp flanking) capture additional regulatory signal, consistent with TF binding cooperativity at adjacent sites.

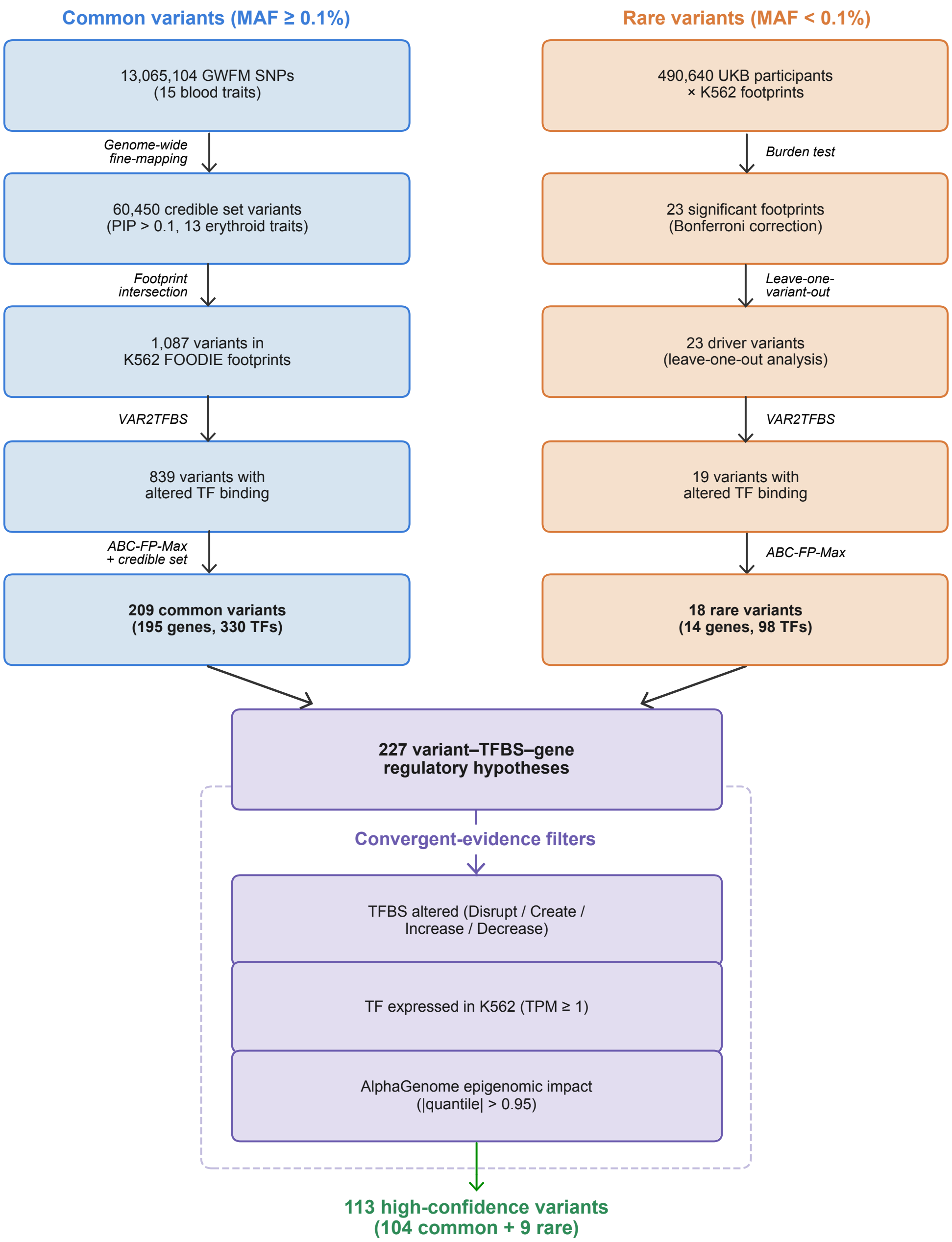


**Supplementary Figure 4.** Multi-layer filtering workflow of the varTFBridge framework. Two parallel pipelines process common variants (left, MAF ≥ 0.1%) and rare variants (right, MAF < 0.1%) through progressive filtering stages. Common variants are filtered from 13,065,104 genome-wide fine-mapped SNPs to 209 credible variants with predicted TF binding changes and target gene assignments. Rare variants are filtered from burden-test-significant FOODIE footprints to 18 driver variants with TF binding effects and gene links. The two branches merge into 227 variant–TFBS–gene regulatory hypotheses, which are further prioritised by three convergent-evidence filters: (1) predicted TFBS alteration, (2) TF expression in K562 (TPM ≥ 1), and (3) AlphaGenome epigenomic impact (|quantile| > 0.95), yielding 113 high-confidence variants (104 common, 9 rare). Related to the section 'A high-confidence resource of variant–TFBS–gene regulatory hypotheses for erythroid traits'.

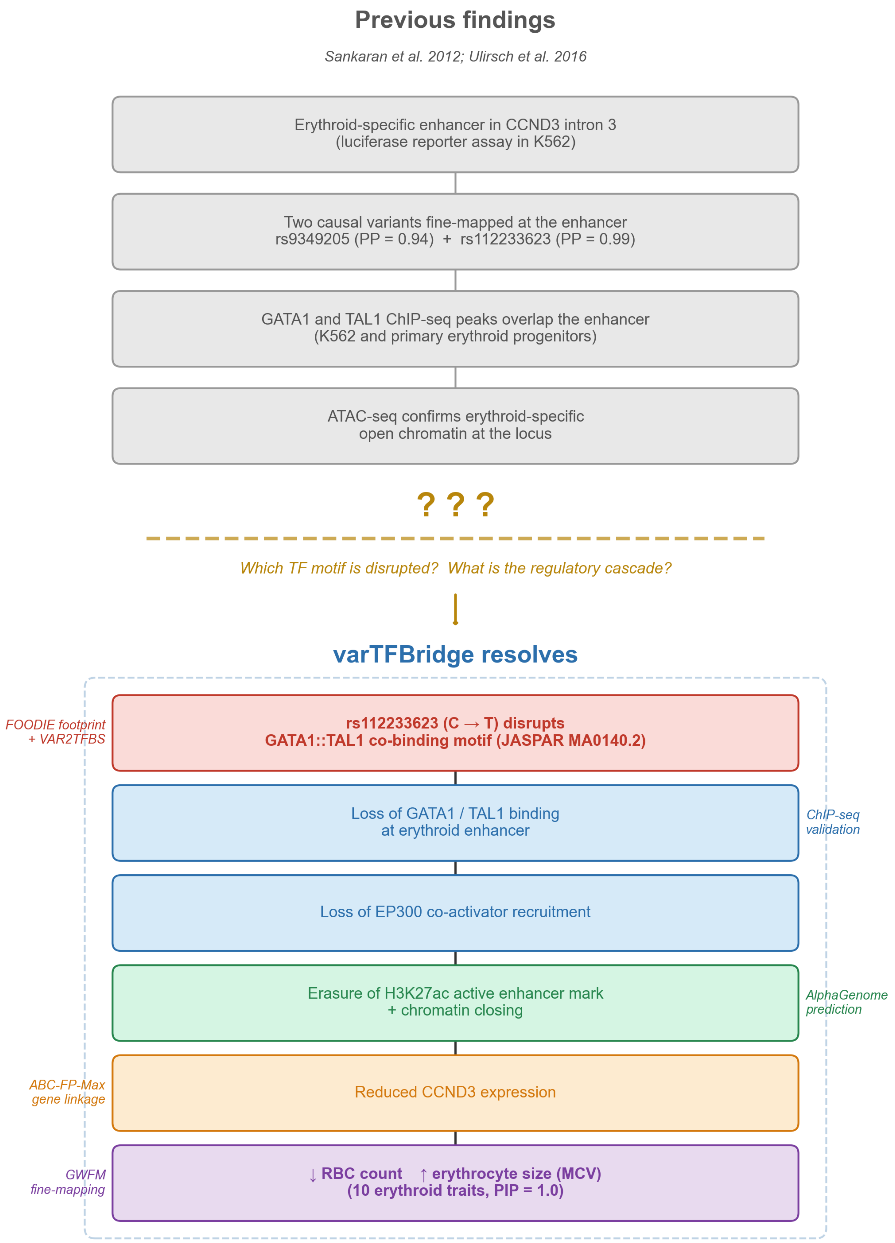

Previous findings
Sankaran et al. 2012; Ulirsch et al. 2016
Erythroid-specific enhancer in CCND3 intron 3
(luciferase reporter assay in K562)
Two causal variants fine-mapped at the enhancer
rs9349205 (PP = 0.94) + rs112233623 (PP = 0.99)
GATA1 and TAL1 ChIP-seq peaks overlap the enhancer
(K562 and primary erythroid progenitors)
ATAC-seq confirms erythroid-specific
open chromatin at the locus
? ? ?
Which TF motif is disrupted? What is the regulatory cascade?
varTFBridge resolves
FOODIE footprint
+ VAR2TFBS
rs112233623 (C → T) disrupts
GATA1::TAL1 co-binding motif (JASPAR MA0140.2)
Loss of GATA1 / TAL1 binding
at erythroid enhancer
ChIP-seq
validation
Loss of EP300 co-activator recruitment
Erasure of H3K27ac active enhancer mark
+ chromatin closing
AlphaGenome
prediction
ABC-FP-Max
gene linkage
Reduced CCND3 expression
GWFM
fine-mapping
↓ RBC count ↑ erythrocyte size (MCV)
(10 erythroid traits, PIP = 1.0)

**Supplementary Figure 5.** Mechanistic resolution of the CCND3 erythroid enhancer locus by varTFBridge. Top (grey): previous findings from Sankaran et al. (2012) and Ulirsch et al. (2016) established that an erythroid-specific enhancer in CCND3 intron 3 harbours two causal variants (rs9349205, PP = 0.94; rs112233623, PP = 0.99) with GATA1 and TAL1 ChIP-seq peaks and erythroid-specific open chromatin, but the specific TF motif disrupted and the downstream regulatory cascade remained unknown. Bottom (coloured): varTFBridge resolves the mechanistic gap by predicting that rs112233623 (C → T) disrupts a GATA1::TAL1 heterodimer co-binding motif (JASPAR MA0140.2), leading to loss of GATA1/TAL1 binding, loss of EP300 co-activator recruitment, erasure of the H3K27ac active enhancer mark, chromatin closing, and reduced CCND3 expression, ultimately resulting in decreased RBC count and increased erythrocyte size (MCV). Side annotations indicate the varTFBridge evidence layer supporting each step: FOODIE footprint detection and VAR2TFBS motif scanning, ChIP-seq validation, AlphaGenome variant effect prediction, ABC-FP-Max gene linkage, and GWFM fine-mapping (PIP = 1.0 across 10 erythroid traits). Related to the section 'varTFBridge predicts the molecular mechanism underlying GATA1/TAL1-dependent CCND3 enhancer regulation'.
