## Supplementary Notes for "Genome-wide maps of transcription factor footprints identify noncoding variants rewiring gene regulatory networks"

**Table of Contents**

Supplementary Note 1. Characterisation of the varTFBridge regulatory resource

Supplementary Note 2. AlphaGenome variant effect predictions for the CCND3 locus

### Supplementary Note 1

**Characterisation of the varTFBridge regulatory resource**

*Related to: “A high-confidence resource of variant–TF–gene regulatory hypotheses for erythroid traits”*

The varTFBridge pipeline produced a resource of 227 variants (209 common, 18 rare) linked to predicted functional TFs and target genes across 13 erythroid traits. Below we describe the composition of this resource in terms of trait representation, TF family involvement, and genomic localisation of linked target genes.

#### Trait distribution

Across the 209 common variants, red cell volume and content traits contributed the largest number of associations: MCV (n = 133), MCH (n = 118), MSCV (n = 96), and MCHC (n = 94), consistent with the erythroid identity of K562. By contrast, reticulocyte traits (RC and RP) contributed the fewest associations (n = 27 and 22, respectively), reflecting higher tissue specificity of reticulocyte regulatory programs.

#### TF distribution

Each variant was linked to an average of 5.7 predicted functional TFs (median = 4; range 1–33), reflecting the dense occupancy of FOODIE footprints by multiple TFs. The most frequently disrupted or created TF binding sites involved members of the GATA (GATA1, GATA2), KLF (KLF1, KLF3, KLF5), and SP (SP1, SP2, SP3) families — all well-established erythroid regulators — as well as broadly acting factors such as CTCF, which anchors chromatin loop boundaries at many loci.

#### Gene and variant localisation

The 227 variants were linked to 207 unique predicted target genes. The majority of variants (131 of 227; 58%) resided within intergenic regions, followed by intronic (79; 35%) and promoter-proximal (17; 7%) locations, indicating that most regulatory effects act through distal enhancer elements rather than proximal promoters.

### Supplementary Note 2

**AlphaGenome variant effect predictions for the CCND3 locus**

*Related to: “varTFBridge predicts the molecular mechanism underlying GATA1::TAL1-dependent CCND3 enhancer regulation”*

AlphaGenome variant effect predictions for K562 provide independent, sequence-based support for the regulatory cascade initiated by GATA1::TAL1 motif disruption at rs112233623 (chr6:41957259, hg38). Below we report the predicted effects on TF binding and downstream chromatin state.

#### Predicted loss of TF binding

The alternative allele is predicted to reduce GATA1 binding (TF ChIP quantile = −0.999) and TAL1 binding (quantile = −1.000), consistent with both the predicted GATA1::TAL1 motif disruption and the observed ChIP-seq occupancy. AlphaGenome further predicts concordant loss of EP300 (quantile = −0.999) — whose binding at this position is experimentally confirmed by ChIP-seq — suggesting that disruption of the GATA1::TAL1 motif leads to loss of EP300 co-activator recruitment and consequently reduced H3K27ac deposition.

#### Predicted chromatin and downstream effects

Consistent with the predicted loss of EP300 recruitment, the model predicts reduced H3K27ac (quantile = −0.999), decreased chromatin accessibility (ATAC-seq quantile = −0.998; DNase-seq quantile = −0.998), and loss of binding for additional erythroid regulators including GATA2 (quantile = −1.000), STAT5A (quantile = −1.000), and SOX6 (quantile = −0.999). All quantile scores exceed the 99.8th percentile of common variant effects, indicating an exceptionally strong predicted regulatory impact.
